# Possible biological control of ash dieback using the parasitic Hymenoscyphus fraxineus mitovirus 2?

**DOI:** 10.1101/2023.03.03.530786

**Authors:** Wajeeha Shamsi, Jana Mittelstrass, Hideki Kondo, Sven Ulrich, Daniel Rigling, Simone Prospero

## Abstract

Invasive fungal diseases represent a major threat to forest ecosystems worldwide. As fungicides are often unfeasible and not a sustainable solution, only a few other control options are available, including biological control. In this context, the use of parasitic mycoviruses as biocontrol agents of fungal pathogens has recently gained particular attention. Since the 1990s, the Asian fungus *Hymenoscyphus fraxineus* has been causing lethal ash dieback across Europe. In the present study, we investigated the biocontrol potential of the mitovirus Hymenoscyphus fraxineus mitovirus 2 (HfMV2) previously identified in Japanese populations of the pathogen. HfMV2 could be successfully introduced via co-culturing into 16 out of 105 virus-free isolates. A virus infection had contrasting effects on fungal growth *in vitro*, from cryptic to detrimental or beneficial. Virus-infected *H. fraxineus* isolates whose growth was reduced by HfMV2 showed a lower virulence on ash (*Fraxinus excelsior*) saplings compared to their isogenic virus-free isolates. The results suggest that mycoviruses exist in the native populations of *H. fraxineus* in Asia that have the potential for biological control of ash dieback in Europe.

## 1. Introduction

About 30% of the world’s land is covered with forests (Keenan et al., 2015) which harbor the richest biodiversity among terrestrial ecosystems and accomplish a broad variety of functions, from providing habitat to wildlife to modulating climate and contributing to improve human living conditions (Linnakoski and Forbes, 2019; Randhir and Erol, 2013). Among the current and future threats to the survival and sustainability of forest ecosystems, invasive pests and diseases play a major role (Fisher et al., 2016). Since the main pathway of the spread of tree pests and pathogens is the trade of live plants, if globalization and international trade intensify, the unintentional movement of potentially dangerous species will most likely continue to increase (Ghelardini et al., 2016).

Successful control of invasive pathogens in forest ecosystems is particularly difficult to achieve because of the specificities of both the hosts and the pathogens (for details, see Prospero et al. (2021)). Given that the application of fungicides is often unfeasible and does not represent a sustainable solution (e.g., risk of development of resistance in the pathogen, possible problems for human and animal health), only a few other options are available, including classical biological control. This implies the introduction of a natural enemy (biological control agent), which will establish a sufficiently large population to control the target pest or pathogen (Kenis et al., 2019). As biocontrol agents, parasitic fungi or bacteria that negatively affect the fitness of their fungal host are usually selected. However, in recent years mycoviruses reducing host virulence have been gaining increasing attention, particularly against fungal pathogens of trees (Ćurković-Perica et al., 2022; Ghabrial and Suzuki, 2009). Following the successful control in Europe of chestnut blight caused by the invasive ascomycete *Cryphonectria parasitica* by a parasitic hypovirus (Cryphonectria hypovirus 1; CHV1) (Anagnostakis, 1982; Heiniger and Rigling, 1994), several studies were undertaken to search for viruses as potential biocontrol agents for other pathogenic fungi. Major invasive fungal pathogens of trees in which mycoviruses have been detected include *Ophiostoma novo-ulmi*, one of the causal agents of Dutch elm disease (Hintz et al., 2013), *Fusarium circinatum*, the causal agent of pine pitch canker (Martínez-Álvarez et al., 2014; Muñoz-Adalia et al., 2016), and *Hymenoscyphus fraxineus*, the causal agent of ash dieback (Čermáková et al., 2017; Schoebel et al., 2014; Shamsi et al., 2022).

Although most tested fungal pathogens were found to be infected with mycoviruses, not all viruses may be suited as biocontrol agents. Besides reducing the virulence of its fungal host (i.e., causing hypovirulence), a virus to be used as a biocontrol agent must fulfil the three following criteria (Rigling et al., 2021): (i) be transmissible to virus-free strains of the host, (ii) override the defense mechanism of the host, and (iii) replicate and propagate across the mycelial network of the host. Even when fulfilling these criteria, a practical hurdle in applying mycoviruses as biocontrol agents may be represented by their inconsistent behavior *in vitro* (agar plates) and *in planta* (trees). For example, although Rosellinia necatrix partitivirus 6-W113 causes severe growth inhibition of the experimental fungal host *Cryphonectria parasitica* on agar medium, it does not reduce fungal virulence toward trees (Suzuki et al., 2021).

Over the last two decades, *H. fraxineus* has caused lethal dieback of common ash (*Fraxinus excelsior*) and narrow-leaved ash (*Fraxinus angustifolia*) across Europe. The fungus is native to East Asia, where it behaves as a saprophytic leaf decomposer (Zhao et al., 2013), and was presumably introduced to Europe in the early 1990s (Kowalski and Holdenrieder, 2009). On susceptible ash species, *H. fraxineus* produces necrotic lesions on leaves, twigs, and stems, eventually leading to wilting and dieback of girdled shoots (Gross et al., 2014). The fungus is heterothallic and reproduces sexually on the ash petioles in the litter once a year. The life cycle is completed during summer when wind-dispersed sexual spores (ascospores) infect new leaves (Kowalski and Holdenrieder, 2009). To date, no efficient control strategy is available for ash dieback. Once established, the disease is difficult to manage since only a small fraction of the ash trees seem to be resistant or tolerant (McKinney et al., 2014; McKinney et al., 2011; Sollars et al., 2017). Regarding the possible use of mycoviruses as biocontrol agents against ash dieback, limited data are available. In the European population of the fungus, Schoebel et al. (2014) described a capsidless single-stranded RNA virus (Hymenoscyphus fraxineus mitovirus 1) that replicates in the mitochondria of the fungal host. However, its high prevalence (up to 90% of the *H. fraxineus* isolates infected) and the lack of clear effects on host virulence (Lygis et al., 2017) seem to strongly limit its potential as a biocontrol agent.

In the present study, we investigated the potential of a second mitovirus (Hymenoscyphus fraxineus mitovirus 2, HfMV2) recently identified in the native population of *H. fraxineus* (Shamsi et al., 2022), to be used as a biocontrol agent against ash dieback. Indeed, specialized and co-evolved enemies (including hyperparasites) are more likely to occur in the native range of a pathogen than in the introduced range. Investigations included the assessment of the effect of HfMV2 on the growth rate of the fungus on agar medium and the fungal virulence toward living ashes.

## 2. Materials and Methods

### 2.1. Fungal isolates

In total, 105 isolates of *H. fraxineus* from two Japanese populations were used, which were screened for mycoviruses in a previous study (Shamsi et al., 2022). One isolate (R1153) was found to be infected with the mitovirus (HfMV2), while the rest of the isolates were HfMV2-free.

### 2.2. Horizontal transmission of HfMV2 to virus-free fungal isolates

To transmit HfMV2 from the virus-infected *H. fraxineus* isolate (donor) to a virus-free isolate (recipient), the paired culture technique was used (Wu et al., 2010a). The donor isolate (R1153) was co-cultured with each of 104 recipient isolates in triplicate on diamalt agar (20 g/L Diamalt, Hefe Schweiz, Stettfurt, Switzerland; 15g/L plant propagation agar, Condalab, Madrid, Spain) (Figure 1A). Plates were incubated for a month in the dark at room temperature. Afterward, agar plugs (5 mm in diameter) taken from three different positions of the paired cultures, i.e., one from the donor isolate and two from the recipient isolate (one in the middle and one from the edge), were sub-cultured to new diamalt agar plates. Virus transmission to the recipient isolate was assessed by one-step reverse transcription polymerase chain reaction (RT-PCR) as described by Urayama et al. (2015) using virus sequence-specific primers (Mt-59-F + Mt-59-R) for HfMV2 (Shamsi et al., 2022).

**Figure 1.**
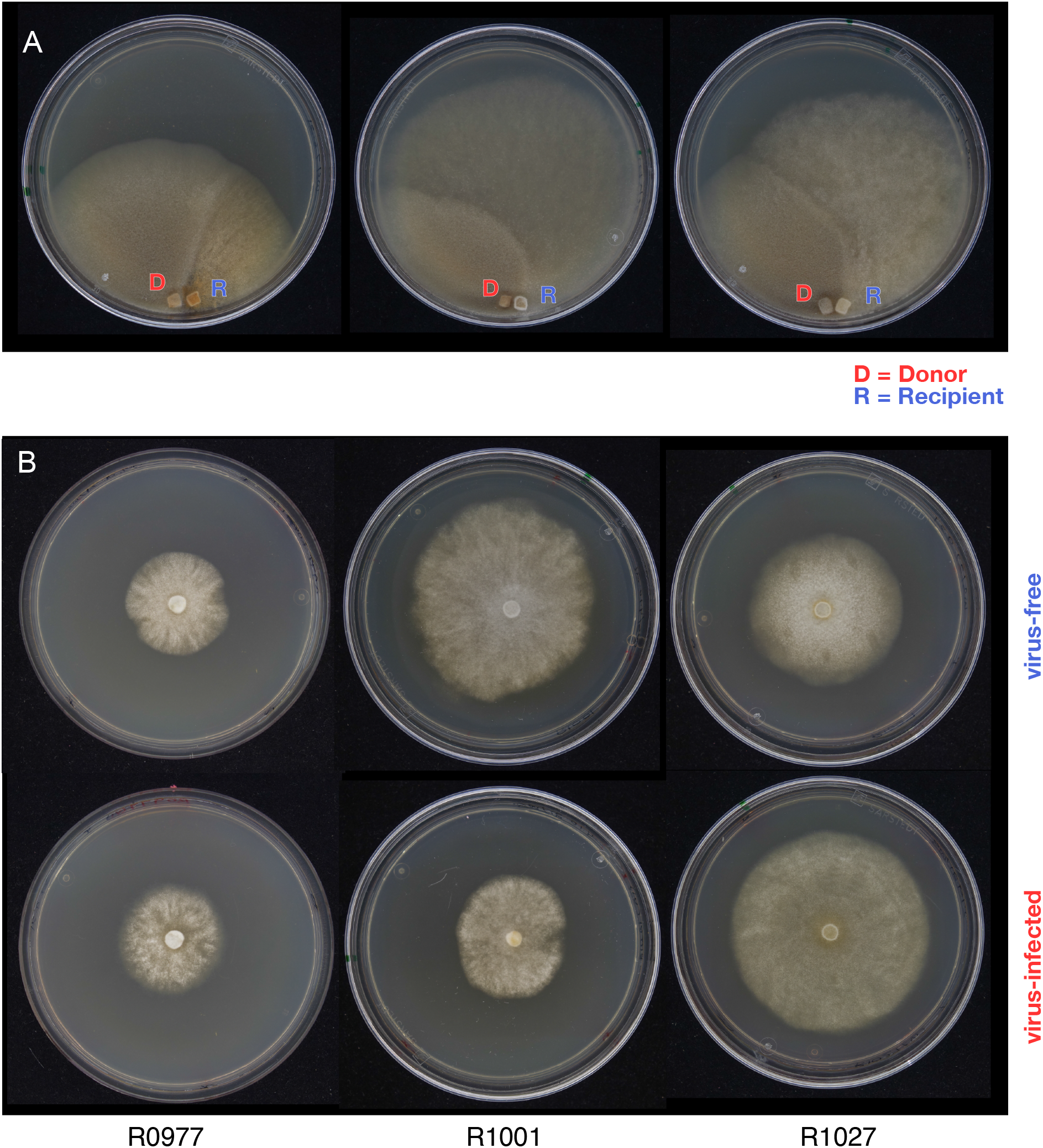
Co-culturing of HfMV2-free and HfMV2-infected isolates of *Hymenoscyphus fraxineus* and effect of HfMV2 on the fungal growth *in vitro*. (A) Representative pairings of HfMV2-free (Recipient; wild-type) and HfMV2-infected (R1153; Donor) *H. fraxineus* isolates after incubation for a month at room temperature. (B) Colony sizes of wild-type HfMV2 and isogenic HfMV2-infected isolates of *H. fraxineus* 15 days post-inoculation on diamalt agar plates. The selected isogenic lines show the three different effects conferred by the virus: R0997→cryptic (no effect), R1001→ detrimental (growth reduction), and R1027→beneficial (growth increase).

Virus-infected isogenic lines of the recipient isolates were obtained by co-culturing the experimentally virus-infected isolates (originally virus-free) with the corresponding virus-free (wild-type) isolate in two rounds.

### 2.3. Microsatellite genotyping

Before pairing, the donor isolate R1153 and all recipient isolates were genotyped at microsatellite loci (Gross et al., 2012a) as described in Burokiene et al. (2015). Three polymorphic microsatellite loci (38, 90, 97; see Supplementary table 1) were selected to discriminate between the virus donor (R1153) and all artificially virus-infected isolates. After pairing, the recipient isolates (wild-type virus-free, but after pairing virus-infected) were transferred to new diamalt plates and subsequently genotyped at the three selected loci to confirm virus transmission into the recipient isolates (Supplementary table 2).

### 2.4. Growth rate comparison

The growth of each virus-free and its corresponding isogenic virus-infected *H. fraxineus* isolate was compared *in vitro* on diamalt agar. Each isolate was inoculated in the middle of a Petri plate (9 cm in diameter) by taking an agar plug (5 mm in diameter) from the margins of a freshly growing culture. Three replicates of each isolate were produced. The colony diameter was measured with a millimeter ruler every fifth day for 15 days (total of three measurements) along two orthogonal lines previously drawn on the plates (after this period, several Petri plates were completely colonized by the fungus). At each measurement, the colony size was calculated as the mean of the two diameters.

### 2.5. Leaf inoculations

Six different pairs of *H. fraxineus* isolates (wild-type virus-free - VF and isogenic virus-infected - VI: R0933, R0982, R1002, R0958, R0969, R0971) were used for the leaf inoculation experiment. The growth of these isolates *in vitro* was shown to be reduced by an infection by HfMV2 (see Results). As tree hosts, 16 healthy three-year-old common ash (*Fraxinus excelsior*) saplings obtained from a Swiss nursery were used. The susceptibility of these saplings to *H. fraxineus* was not known.

Inoculations were performed in the biosafety level 3 greenhouse facility at WSL as described by Orton et al. (2019). Using a scalpel, approximately 1 cm^2^ of the agar with mycelium (3-4 mm thickness) was excised from a 7-day-old *H. fraxineus* culture and placed on a wound previously produced on the epidermis of a healthy rachis, between the second and third pair of leaflets. At the lower side of the wound, the epidermis was left attached to the rachis. The fungal inoculum was covered by the epidermis and parafilm was wrapped around it as protection against desiccation. Each sapling was inoculated with two different pairs of virus-infected and virus-free isolates. Five replicates of each pair of isolates were inoculated on different trees. Mock inoculations with agar were used as a negative control.

### 2.6. Data analysis

The results of the two inoculation experiments (*in vitro* and *in planta*) were visualized and analyzed using the software *R* (version 4.1.0) (Team, 2012). For each isolate included in the *in vitro* growth assay, linear models (LMs) were used to test the effect of HfMV2 on the fungal growth at five, 10, and 15 days after inoculation. To assess the effect of the fungal isolate, virus presence, and their interaction on the probability to develop an infection, binomial generalized linear models (GLMs) were fit to the data using the function *glm*. The factor host tree was included in the models as a random effect.

## 3. Results

### 3.1. Effects of HfMV2 on the growth of *H. fraxineus in vitro*

HfMV2 transmission from a donor *H. fraxineus* isolate (R1153) to virus-free recipient isolates was successful in 16 out of 104 (15.4%) paired cultures. RT-PCR detected the virus in all mycelial samples taken from the 16 initially virus-free isolates. All the artificially virus-infected isolates showed the expected microsatellite pattern of the recipient isolates, confirming successful virus transmission. (Supplementary table 1). The virus was stable after several subcultures in the recipient isolates.

To assess the effect of HfMV2 on the growth of *H. fraxineus in vitro*, the growth of virus-free and virus-infected isogenic lines was compared (Figure 1B, Figure 2). The sizes of cultures from the virus-free and virus-infected lines increased in a similar manner over the course of 15 days post-inoculation (Figure 2A). We, thus, compared the final culture sizes (after 15 days) of the virus-free and virus-infected lines within separate isolates (Figure 2B, Supplementary table 3). In ten out of 16 isolate pairs, HfMV2 significantly affected the size of the fungal colony. In eight of these pairs, the virus-infected isolates showed smaller culture sizes than their corresponding isogenic virus-free isolates. In seven pairs, no significant difference in the size of the virus-infected and virus-free isolates was detected at the final measurement. Noteworthy, isolate R1027 showed increased culture sizes in the virus-infected isolate compared to the virus-free isolate.

**Figure 2.**
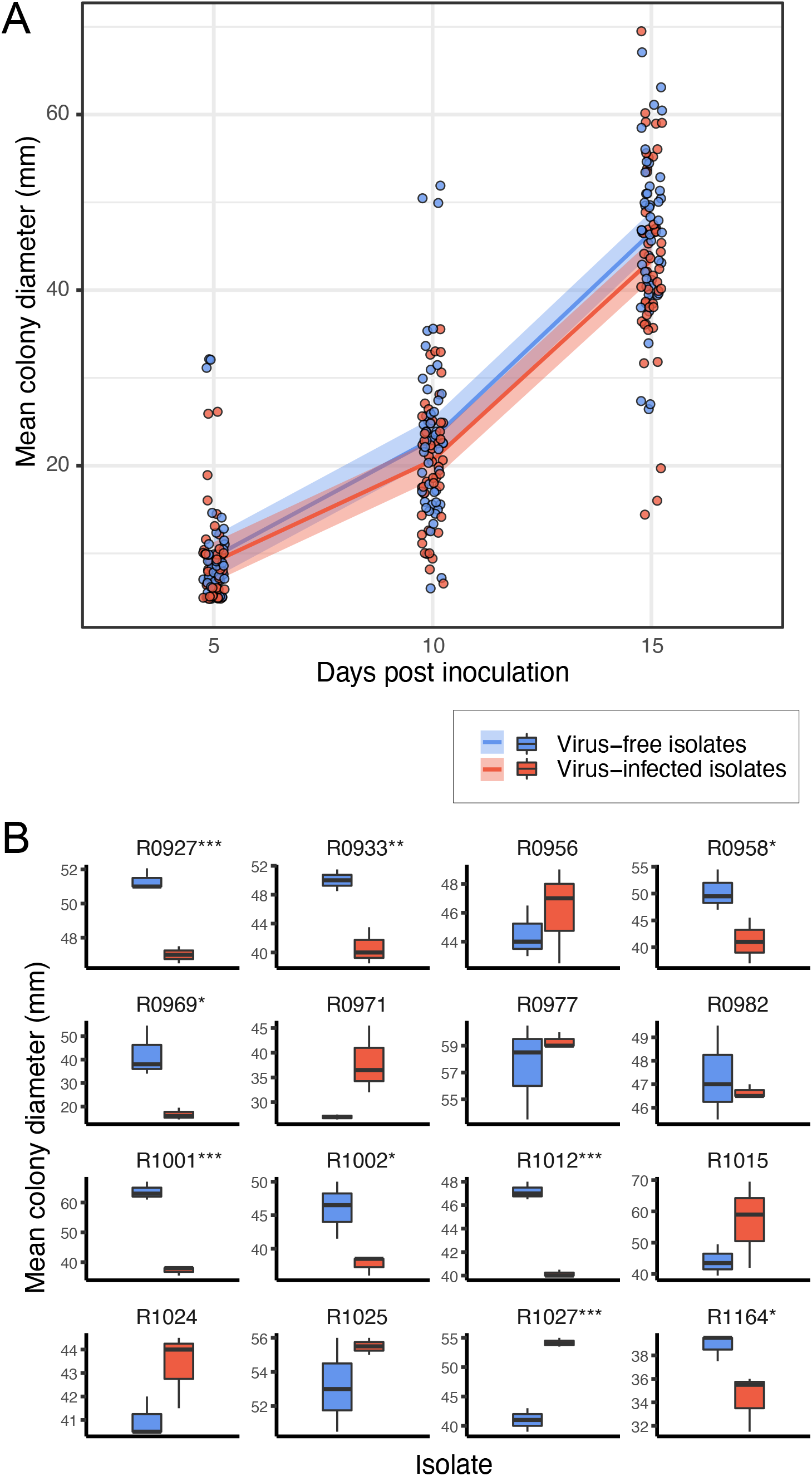
Growth of HfMV2-free and HfMV2-infected *Hymenoscyphus fraxineus* isolates *in vitro* over time. (A) The mean colony diameter of all isolate replicates is shown over the course of 15 days post-inoculation. A ‘loess’ regression line was fit for each of the two groups (virus-free and virus-infected isolates). **(B)** The colony sizes of virus-free and virus-infected strains at 15 days post-inoculation are shown for each isolate separately. Statistical significance displayed in the plots correspond to *P*-values from F-tests evaluating the effect of the virus within each isolate (n=3 for each level); \*\*\**P*<0.001, \*\**P*<0.01, \**P*<0.05.

### 3.2. Effect of HfMV2 on fungal virulence

At the end of the *in planta* experiment (i.e., 10 weeks post-inoculation), 16 out of 31 inoculated rachises (51.6%) showed a visible lesion and wilting leaves distal to it (Figure 3A), indicating a successful infection by *H. fraxineus:* 10 rachises were inoculated with a virus-free isolate, whereas six rachises were inoculated with a virus-infected isolate. All negative controls (mock inoculations) showed no symptoms. In five ash saplings (# 2, 4, 11, 12, 14) neither the virus-free nor the virus-infected isolate caused symptoms, while in 9 saplings (# 1, 3, 5, 6, 7, 8, 9, 13, 15) symptoms were induced only by one of the inoculated pair of isolates. Only in two saplings (# 10 and 16), lesions and wilting leaves appeared on all inoculated rachises.

**Figure 3.**
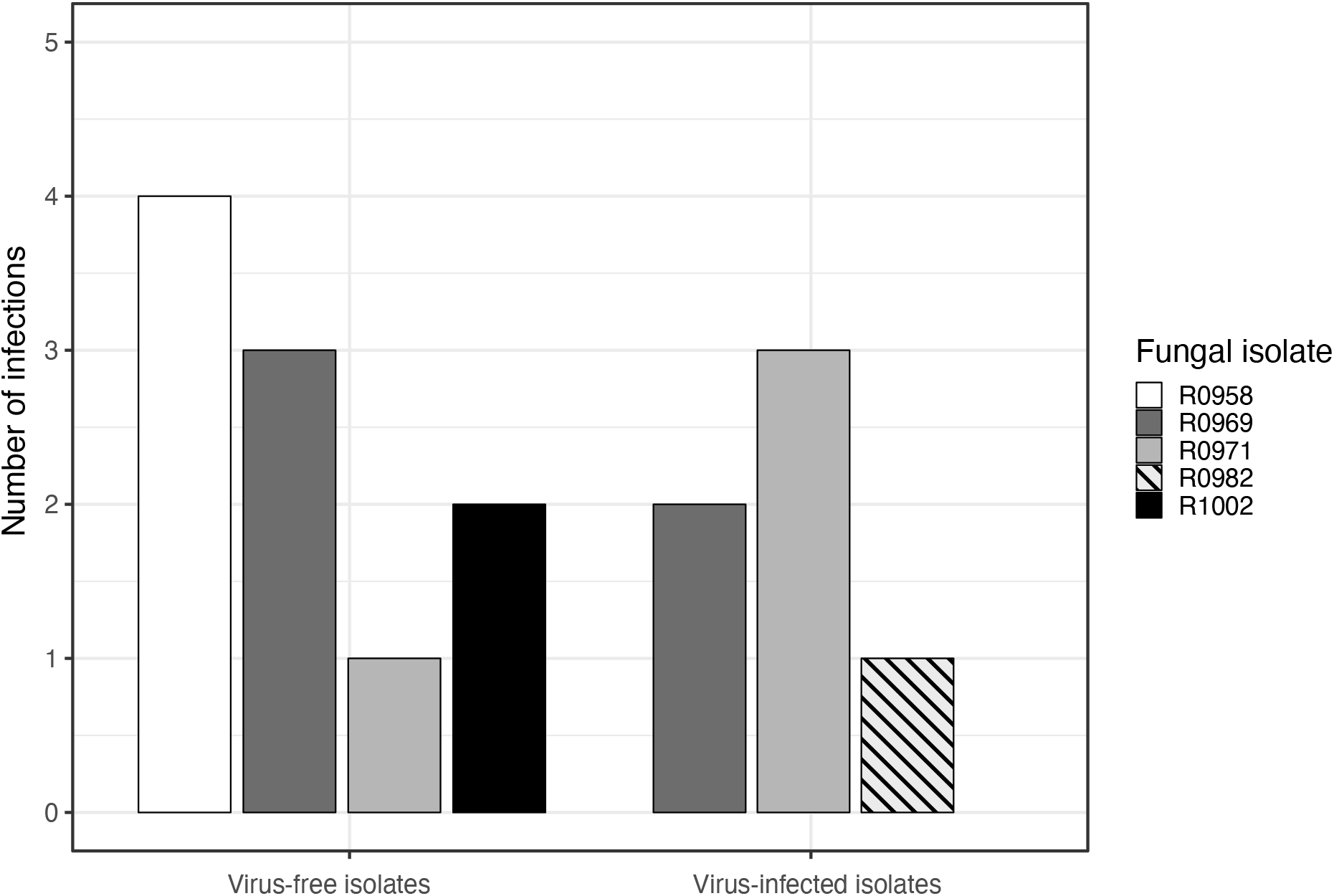
Total number of rachis inoculations with HfMV2-free and HfMV2-infected *Hymenoscyphus fraxineus* isolates that resulted in symptoms (wilting leaves distal to the inoculation point) in the greenhouse experiment. The number of replicates was n=5 for both virus-free and virus-infected isolates, except for isolate R0982 (n=6 for both). Data for the isolate R0933 is not shown, as inoculations with it did not result in any symptoms.

All inoculated *H. fraxineus* isolates (R0982, R1002, R0958, R0969, R0971) except R0933 resulted in at least one infection, either by the wild-type virus-free isolate or by the isogenic virus-infected isolate or both (Figure 3). Overall, more infections were produced by virus-free than virus-infected isolates. However, important differences were observed within pairs of isogenic virus-free and virus-infected isolates. Isolates R0958 and R1002 only caused symptoms when virus-free (4 and 2 infections, respectively), while on the other side isolate R0982 induced a lesion with wilting leaves only when virus-infected. Both with virus-free and virus-infected isolates, first symptoms of an infection (wilting leaves) appeared three weeks after inoculation (Figure 4). The number of infections caused by virus-free isolates rapidly increased until week 8 and then remained constant. The increase in the number of infections caused by virus-infected isolates was lower and continued till the end of the experiment. GLMs revealed a significant effect of the fungal isolate on the probability to develop a leaf infection (Table 1). Although there was no additive effect of fungal isolate and virus, the effect of the interaction between the virus and fungal isolate was also significant (Table 1).

**Figure 4.**
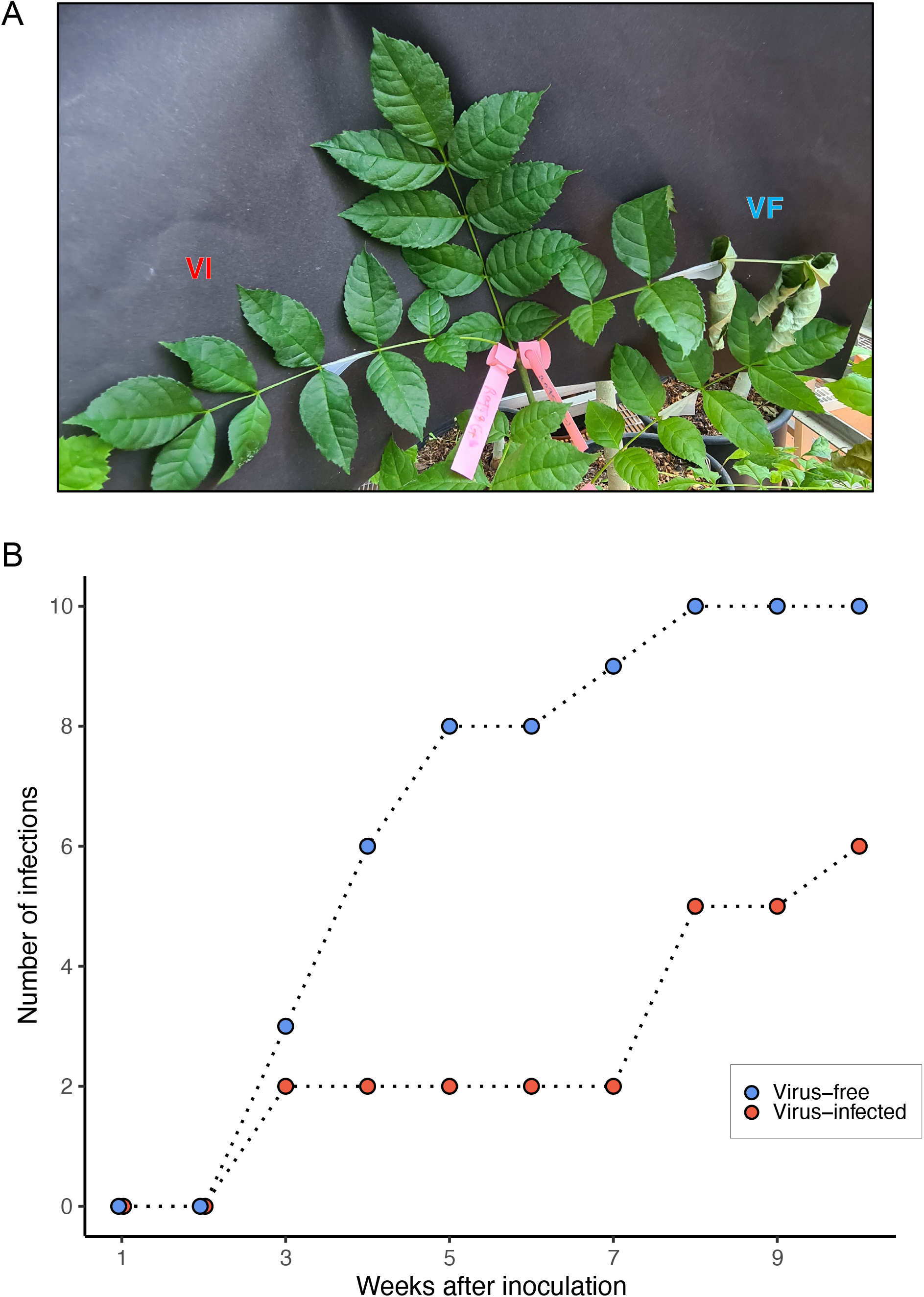
Inoculation of rachises of *Fraxinus excelsior* saplings with selected isolates of *Hymenoscyphus fraxineus* in the greenhouse experiment. **(A)** Representative inoculation (with isolate R0958) showing dieback symptoms in the leaflet inoculated with a virus-free isolate (VF), but not in the leaflet inoculated with a virus-infected isolate (VI). **(B)** Cumulative number of infected (i.e., with wilting leaves distal to the inoculation point) leaflets over the course of the experiment (10 weeks). Blue dots: virus-free *H. fraxineus* isolates, red dots: virus-infected *H. fraxineus* isolates.

**Table 1.** Influence of the fungal isolate, virus presence, and their interaction, on the development of an infection tested with binomial generalized linear models (GLMs).

| Explanative variables | Resid. Df<sup>1</sup> | Degrees of freedom | *P*-value<sup>2</sup> |
| --- | --- | --- | --- |
| infection ~ 1 | 61 | NA<sup>3</sup> | NA |
| infection ~ tree | 60 | 1 | 0.368 |
| infection ~ tree + fungal_isolate | 55 | 5 | 0.030* |
| infection ~ tree + fungal_isolate + virus | 54 | 1 | 0.199 |
| infection ~ tree + fungal_isolate * virus | 49 | 5 | 0.018* |

<sup>1</sup>Resid. Df = residual degrees of freedom.
<sup>2</sup>\* = < 0.05.
<sup>3</sup>NA = not applicable.

## 3.3. Discussion

In this study, we investigated the effect of the mycovirus HfMV2 on the fitness of the fungal host *H. fraxineus* and its horizontal transmissibility within the fungal population. HfMV2, which naturally occurs in the native *H. fraxineus* population in Japan (Shamsi et al. 2022), might represent a hope for the biological control of the devastating ash dieback in Europe. However, to have a future as a biological control agent, a parasitic mycovirus needs to fulfil several criteria (Rigling et al., 2021), among which, the basic one is to reduce the virulence of its fungal host. To assess the outcome of the fungus-virus interaction, it is essential to produce pairs of isogenic fungal strains that differ only in the presence or absence of the target mycovirus. Although introducing a virus into a virus-free fungal strain or curing an originally infected fungal strain is not always easy, great advancements have been made in the past decades (Ghabrial et al., 2015; Kondo et al., 2013; Pearson et al., 2009; Shahi et al., 2019). Here, we used the paired culture technique method, which was already successfully applied in several previous studies (Khalifa et al., 2022; Khalifa and Pearson, 2013; Wu et al., 2007). By growing virus-free and virus-infected fungal cultures in proximity on an agar medium, the hyphae of the two cultures may fuse (anastomose), thereby allowing the transmission of cytoplasmic elements, like viruses.

An infection by HfMV2 had three different effects on the growth of *H. fraxineus in vitro*, namely detrimental (growth reduction), beneficial (growth increase), and cryptic (no change). Variable effects of mitoviruses on their fungal hosts were also reported in previous studies (Flores-Pacheco et al., 2017; Romeralo Tapia et al., 2012; Wu et al., 2007). In particular, Hyder et al. (2013) observed contrasting results when they compared the effects of two viruses on several virus-free isolates of *Heterobasidion* wood decay fungi. According to the authors, the effects of a single virus strain on the fitness of a fungal isolate may vary according to environmental and ecological conditions. Since in our experiment all *H. fraxineus* isolates were incubated under the same conditions, we cannot conclude about the influence of environmental conditions on the outcome of the virus-fungus interaction. Nevertheless, our study confirms the crucial role of the genetic background of the fungal host. In general, in filamentous fungi, RNA silencing is responsible for fungal antiviral defense responses at the cellular level (Nuss, 2011), but the counter-defense mechanism of mycoviruses against RNA silencing is largely unknown. It has been reported that some mycoviruses can suppress the RNA silencing pathway (Wu et al., 2010b). The host genes involved in RNA silencing have been studied in several fungi, including *C. parasitica* (Nuss, 2011), *S. sclerotiorum (Mochama et al., 2018; Neupane et al., 2019*) and *R. necatrix* (Yaegashi et al., 2013). The variable effects induced by HfMV2 on the growth of different *H. fraxineus* isolates may be due to intraspecific differences in the RNA silencing machinery. Zhang et al. (2012) showed a complex interaction between the host RNA silencing antiviral defense and the silencing suppressers of mycoviruses leading to different symptom induction profiles in *C. parasitica* strain EP155 (wild type virus-free) and Δ*dcl-2* (*dicer like 2* mutant). Noteworthy, no phenotypic changes were induced by Cryphonectria parasitica mitovirus 1 on the two *C. parasitica* strains (Shahi et al., 2019). No differences were also observed in CpMV1 genomic RNA accumulation between the wild type and mutant strain. This suggests that CpMV1 may avoid host RNA silencing, which is consistent with its mitochondrial replication (Shahi et al., 2019). Specific analysis of the genes involved in RNA silencing in *H. fraxineus* and their correlation with the RNA silencing suppressors of HfMV2 would help to explain the variable effects of HfMV2 on fungal growth.

Virus-infected *H. fraxineus* isolates whose growth *in vitro* was reduced by HfMV2 showed a lower virulence compared to their isogenic virus-free isolates. Considerable differences in virulence were also observed among virus-free isolates, which may reflect the variation in virulence present in native populations of *H. fraxineus*. Regarding tree response to inoculations, three different situations were recorded: 1) none of the inoculated rachises in a sapling showed infection, 2) some of the inoculated rachises showed infection, or 3) all the inoculations in a sapling resulted in infection. These results indirectly confirm the existence of varying degrees of susceptibility and tolerance to *H. fraxineus* in the European ash population, as previously revealed by specific inoculation experiments (Lobo et al., 2015; McKinney et al., 2012; McKinney et al., 2011; Pliura et al., 2011; Sollars et al., 2017; Stener, 2013).

In addition to causing hypovirulence, a mycovirus needs to infect a large proportion of the pathogen’s population to be useful as a biocontrol agent. Horizontal transmissibility of HfMV2 in the *H. fraxineus* population seems to be quite low since, in paired cultures, only about 15% of the virus-free isolates became virus-infected. Limited data are available about horizontal transmission rates of parasitic mycoviruses in fungal populations. Yang et al. (2018) could artificially infect all tested (n=25) virus-free strains of the plant pathogen *Sclerotinia minor* with the Sclerotinia minor endornavirus 1 using the paired culture method. By co-culturing virus-free and virus-infected isolates, Flores-Pacheco et al. (2017) were able to transmit the Fusarium circinatum mitovirus 1 and 2-2 to all seven virus-free isolates of the pine pitch canker pathogen *Fusarium circinatum*. A major barrier to the successful horizontal transmission of mycoviruses between fungal isolates is represented by vegetative incompatibility, which prevents the spread of suppressive cytoplasmic determinants (Caten, 1972). This phenomenon of non-self-allorecognition is genetically controlled by heterokaryon genes (*het*) or vegetative incompatibility genes (*vic*) (Leslie, 1993). Even though the genetics of the vegetative incompatibility system in *H. fraxineus* is still unknown, a study conducted in the UK showed that 96% of isolate pairings were incompatible (Orton et al., 2018). A similar situation is most likely to occur also in the native populations of the fungus and can be explained by the obligatory outcrossing sexual system of *H. fraxineus*.

Besides contact between mycelia, viruses can also spread in a fungal population through spores, so-called vertical transmission (Nuss, 2005). In some fungal pathogens like *C. parasitica* mycoviruses can be transmitted only by asexual spores, whereas in other species like *H. annosum* sexual spores can also be virus-infected (Ding et al., 2007; Ihrmark et al., 2004; Schoebel et al., 2017). In *H. fraxineus*, asexual conidia seem not to play a role in the spread of the fungus since their function is to act as spermatia during ascospore formation (Gross et al., 2012b). Thus, for a mycovirus to become highly prevalent in the *H. fraxineus* population must be transmissible to the sexual ascospores. To determine whether HfMV2 may spread by ascospores, we tried to artificially produce sexual fruiting bodies (apothecia) and ascospores by crossing virus-infected isolates of the opposite mating type. Unfortunately, our trial was not successful because of technical issues with the incubator. Nevertheless, the finding that another mitovirus (HfMV1), detected in the European population of *H. fraxineus*, is transmitted at high frequency into ascospores (see table 3 in Schoebel et al. 2017) suggest, that such mitoviruses are commonly transmitted into sexual spores.

Our study revealed that the mitovirus HfMV2 previously detected in the Japanese population of *H. fraxineus* can reduce the virulence of at least some isolates of its fungal host. Regarding the possible use of HfMV2 as biological control agent for ash dieback in Europe, several aspects still need further investigation. First, the transmissibility of the mitovirus to the ascospores, which are the main dispersal and infection propagules of *H. fraxineus*. Second, the possibility to artificially infect European isolates of the pathogen with HfMV2 since the present study was performed with Japanese isolates. Besides co-culture technique, protoplast fusion could also be utilized for the artificial introduction of HfMV2 into virus-free isolates (Shahi et al. (2019). Eventually, to enhance HfMV2 transmission across different vegetative compatibility types of *H. fraxineus* a super donor strain could be produced, as done in the chestnut blight fungus *C. parasitica* (Stauder et al., 2019).

## Acknowledgments

This study was conducted in the frame of the HOMED project (http://homed-project.eu/). which received funding from the European Union’s Horizon 2020 research and innovation program under grant agreement No. 771271. S.P. also received financial support with a grant from the Swiss Federal Office for the Environment FOEN.

