## Supplementary data for "Possible biological control of ash dieback using the parasitic Hymenoscyphus fraxineus mitovirus 2?"

Simone Prospero

Address:

Swiss Federal Institute for Forest, Snow and Landscape Research WSL,

Zuercherstrasse 111, 8903 Birmensdorf, Switzerland.

### Supplementary Tables

**Supplementary Table 1.** Microsatellite loci used to genotype *H. fraxineus* isolates.

| Locus name | Abbreviation | Primer concentration | Dye | Allele sizes (bp) | Number of alleles |
|---|---|---|---|---|---|
| mHp_111990 | 90 | 0.25 µM | HEX | 213, 216 | 2 |
| mHp_103438 | 38 | 0.15 µM | ATTO 550 | 231, 234, 237 | 3 |
| mHp_080497 | 97 | 0.30 µM | FAM | 245, 254 | 2 |

**Supplementary Table 2**. Genotyping of the donor and recipient *H. fraxineus* strains used for the horizontal virus transmission. Each unique allele was additionally symbolized by coloured shapes.

| | sample ID | Allele size 90 | Allele size 38 | Allele size 97 | Genotype |
|---|---|---|---|---|---|
| Donor strain | R1153 | 207 | 228 | 245 | |
| Recipient strains | R1027 | 207 | 243 | 245 | |
| | R0977 | 210 | 225 | 245 | |
| | R1015 | 222 | 231 | 245 | |
| | R1164 | 213 | 231 | 245 | |
| | R0971 | 219 | 237 | 245 | |
| | R1025 | 242 | 237 | 245 | |
| | R0956 | 213 | 231 | 245 | |
| | R1024 | 213 | 237 | 245 | |
| | R0982 | 207 | 237 | 245 | |
| | R0958 | 213 | 237 | 248 | |
| | R1002 | 219 | 233 | 248 | |
| | R0933 | 210 | 228 | 245 | |
| | R0969 | 213 | 228 | 248 | |
| | R0927 | 213 | 233 | 248 | |
| | R1001 | 204 | 228 | ? | |
| | R1012 | 213 | 225 | 245 | |

**Supplementary Table 3.** The results of LMs testing the effect of the virus presence on the average size of cultures of each isolate. The formula ‘avg_diam ~ 1 + type_virus’ was used for the data of each separate isolate after 15 days of culture growth.

| Isolate | Days post-infection | *P*-value F-test | Coefficient | R-squared | Statistical significance |
|---|---|---|---|---|---|
| R0927 | 15 | 0.0006 | -4.333 | 0.960 | *** |
| R0933 | 15 | 0.0055 | -9.333 | 0.881 | ** |
| R0956 | 15 | 0.4883 | 1.667 | 0.127 | |
| R0958 | 15 | 0.0499 | -9.167 | 0.659 | * |
| R0969 | 15 | 0.0167 | -25.500 | 0.796 | * |
| R0971 | 15 | 0.0506 | 11.000 | 0.656 | |
| R0977 | 15 | 0.4336 | 1.833 | 0.159 | |
| R0982 | 15 | 0.6018 | -0.667 | 0.074 | |
| R1001 | 15 | 0.0002 | -26.500 | 0.979 | *** |
| R1002 | 15 | 0.0329 | -8.333 | 0.719 | * |
| R1012 | 15 | 0.0001 | -7.000 | 0.982 | *** |
| R1015 | 15 | 0.2114 | 12.667 | 0.356 | |
| R1024 | 15 | 0.0913 | 2.333 | 0.551 | |
| R1025 | 15 | 0.2222 | 2.333 | 0.343 | |
| R1027 | 15 | 0.0004 | 13.167 | 0.966 | *** |
| R1164 | 15 | 0.0458 | -4.500 | 0.672 | * |
